# The key parameters that govern translation efficiency

**DOI:** 10.1101/440693

**Authors:** Dan D. Erdmann-Pham, Khanh Dao Duc, Yun S. Song

## Abstract

Translation of mRNA into protein is a fundamental yet complex biological process with multiple factors that can potentially affect its efficiency. In particular, different genes can have quite different initiation rates, while site-specific elongation rates can vary substantially along a given transcript. Here, we analyze a stochastic model of translation dynamics to identify the key parameters that govern the overall rate of protein synthesis and the efficiency of ribosome usage. The mathematical model we study is an interacting particle system that generalizes the Totally Asymmetric Simple Exclusion Process (TASEP), where particles correspond to ribosomes. While the TASEP and its variants have been studied for the past several decades through simulations and mean field approximations, a general analytic solution has remained challenging to obtain. By analyzing the so-called hydrodynamic limit, we here obtain exact closed-form expressions for stationary currents and particle densities that agree well with Monte Carlo simulations. In addition, we provide a complete characterization of phase transitions in the system. Surprisingly, phase boundaries depend on only four parameters: the particle size, and the first, last and minimum particle jump rates. Relating these theoretical results to translation, we formulate four design principles that detail how to tune these parameters to optimize translation efficiency in terms of protein production rate and resource usage. We then analyze ribosome profiling data of *S. cerevisiae* and demonstrate that its translation system is generally efficient, consistent with the design principles we found. We discuss implications of our findings on evolutionary constraints and codon usage bias.

## 1 Introduction

Being a major determinant of gene expression and protein abundance levels [1, 2], translation of mRNA into polypeptides is one of the most fundamental biological processes underlying life. The extent to which this process is regulated and shaped by the sequence landscape has been widely studied over the past decades [3, 4, 5], revealing many intricate mechanisms that may affect translation dynamics. From a more global perspective, however, it has been challenging to integrate these findings to elucidate the key factors that govern translation efficiency. Indeed, translation is a complex process that depends on many parameters, including the initiation rate, site-specific elongation rates (which can vary substantially along a given transcript), and the termination rate. How does the overall rate of protein synthesis depend on these parameters? To make the problem more concrete, suppose that the goal is to achieve the fastest rate of protein production while minimizing the cost. Would choosing the “fastest” synonymous codon at each site do the job? If the local elongation rate changes at a particular site, would it necessarily affect the overall rate of protein synthesis? If not, then which parameters actually matter? Aside from achieving a desired protein production rate, how does a translation system make efficient use of available resources, particularly the ribosomes?

In this article, we develop a theoretical tool to answer the above questions. Our work hinges on analyzing a mathematical model that describes the traffic of ribosomes, which mediate translation by moving along the mRNA transcript. Beginning with MacDonald *et al.* [6], most mechanistic studies of translation dynamics have been based on the so-called Totally Asymmetric Simple Exclusion Process (TASEP), a probabilistic model that explicitly describes the flow of particles along a lattice [7, 8]. As a classical model of transport phenomena in non-equilibrium, the TASEP has attracted wide interest from mathematicians and physicists [9]. To describe translation realistically, however, a generalized version of the model needs to be employed, taking into account the extended size of the ribosome and the heterogeneity of the elongation rate along the transcript. Under such general conditions, critical questions have hitherto remained open; in particular, identifying the parameters most crucial to the current and particle density has proven elusive.

Here we carry out a theoretical analysis of a generalized version of the TASEP and obtain analytic results that provide new insights into translation dynamics. Our approach is to study the process in a continuum limit called the hydrodynamic limit, which leads to a general PDE satisfied by the density of particles. Upon solving this PDE, we obtain exact closed-form expressions for stationary currents and particle densities that agree very well with Monte Carlo simulations of the original TASEP model. Furthermore, we provide a complete characterization of phase transitions in the system. These results allow us to identify the key parameters that govern translation dynamics, and to formulate a set of specific design principles for optimizing translation efficiency in terms of protein production rate and resource usage. Using experimental ribosome profiling data of *S. cerevisiae*, we show that the translation system of this organism is generally efficient according to the design principles we found.

## 2 Theoretical results on the inhomogeneous *ℓ*-TASEP

### Model description

At a high level, translation of mRNA involves three types of movement of the ribosome, as illustrated in Figure 1a: 1) Initiation - a small ribosomal subunit enters the open reading frame so that its A-site is positioned at the second codon and then a large ribosomal subunit binds with the small subunit. 2) Elongation - the nascent peptide chain gets elongated by one amino acid and the ribosome moves forward by one codon. 3) Termination - the ribosome with its A-site at the stop codon unbinds from the transcript. An important point to note is that more than one ribosome can translate the same mRNA transcript simultaneously, so the movement of a ribosome can be obstructed by another ribosome in front, similar to what happens in a traffic flow on a one-lane road. Such interaction is what makes the dynamics difficult to analyze.

**Figure 1.**
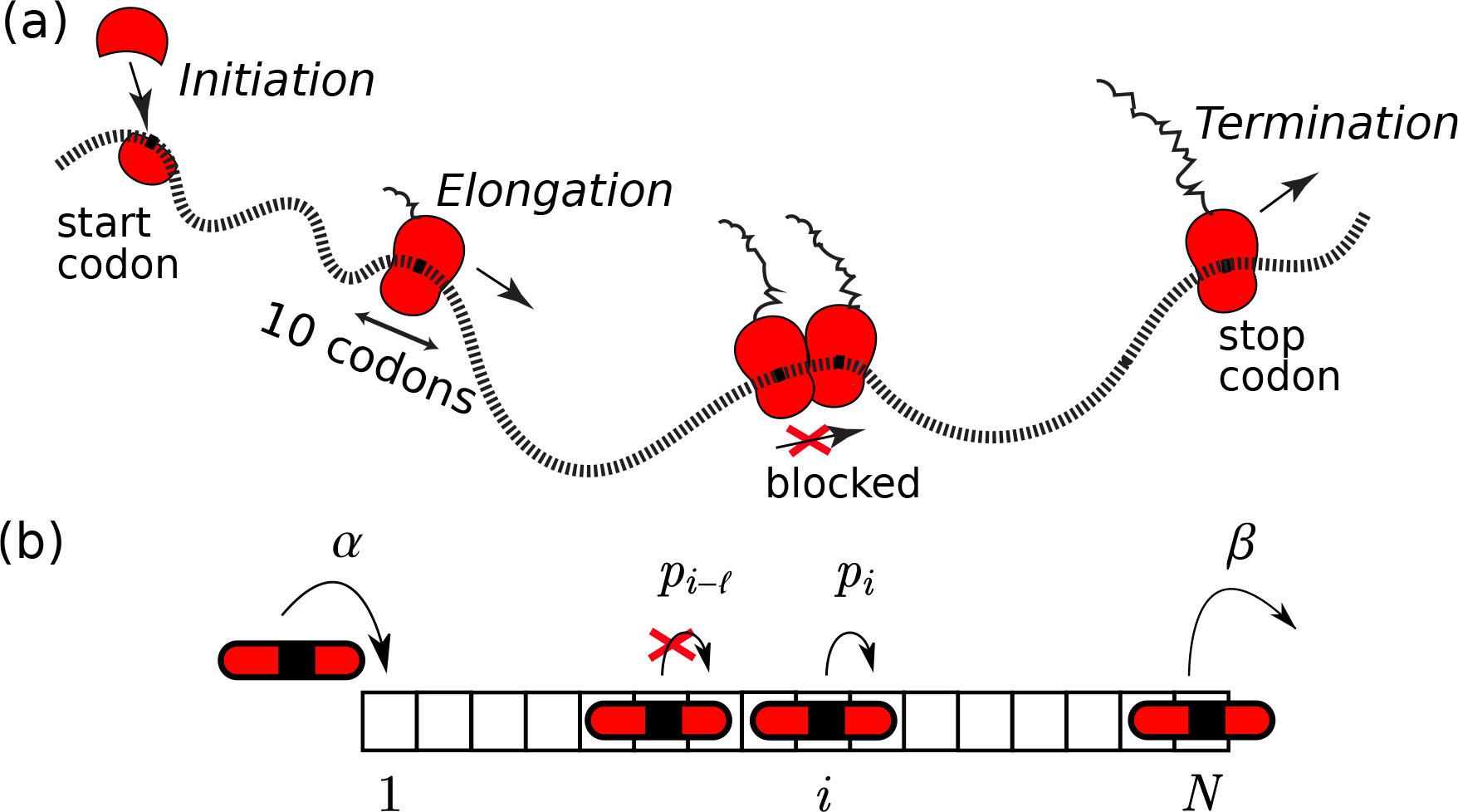
Illustration of the translation process and the inhomogeneous *ℓ*-TASEP with open boundaries. (a) Ribosomes initiate translation at the mRNA 5′ end, elongate the polypeptide by decoding one codon at a time, and eventually terminate the process by detaching from the transcript. **(b)** Particles (of size *ℓ* = 3 here) enter the lattice at rate *α* and a particle at position *i* (here defined by the position of the midpoint of the particle) moves one site to the right at rate *p_i_*, provided that the move is not blocked by another particle in front.

We model the flow of ribosomes on mRNA using a generalized TASEP, called the inhomogeneous *ℓ*-TASEP, on a one-dimensional lattice with *N* sites (see Figure 1b). In this process, each particle is of a fixed size *ℓ* ∈ ℕ and is assigned a common reference point (e.g., the midpoint in the example illustrated in Figure 1b). The position of a particle is defined as the location of its reference point on the lattice. A configuration of particles is denoted by the vector **τ** =(*τ*_1_, …, τ_*N*_), where τ_*i*_ = 1 if the *i*^th^ site is occupied by a particle reference point and τ_*i*_ = 0 otherwise. The jump rate at site *i* of the lattice is denoted by *p_i_* > 0. During every infinitesimal time interval *dt*, each particle located at position *i* ∈ {1, …, *N* - 1} has probability *p*_*i*_*dt* of jumping exactly one site to the right, provided that the next *ℓ* sites are empty; particles at positions between *N* - *ℓ* + 1 and *N*, inclusive, never get obstructed. Additionally, a new particle enters site 1 with probability *αdt* if τ_*i*_ = 0 for all *i* = 1, …, *ℓ*. If *τ_N_* = 1, the particle at site *N* exits the lattice with probability *βdt*. The parameter *α* is called the entrance (or initiation) rate, while *β* is call the exit (or termination) rate.

### The hydrodynamic limit

The key quantities of interest are the stationary probability 〈τ_*i*_〉 of any individual site *i* being occupied or not, and the current (or flux) *J* of particles in the system. In the corresponding translation process, these quantities reflect the local ribosomal density and the protein production rate, respectively.

In the special case of the homogeneous 1-TASEP (*p*_*i*_ = *p* for all *i* and *ℓ* = 1), the stationary distribution of the process decomposes into matrix product states, which can be treated analytically [10]. Unfortunately, in the general case this approach is intractable, necessitating alternative methods such as the hydrodynamic limit. When *ℓ* > 1, deriving the hydrodynamic limit is not straightforward, however, as the process does not possess stationary product measures [11]. To tackle this problem, we mapped the *ℓ*-TASEP to another interacting particle system called the zero range process (ZRP), whose hydrodynamic limit can be derived from the associated master equation. More precisely, we obtained the hydrodynamic limit through Eulerian scaling of time and space by a factor *a* = *N*^-1^, and by following its dynamics on scale *x* such that 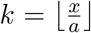, for 1 < *k* < *N* [12]. Implementing this limiting procedure for the ZRP and mapping it back to the inhomogeneous *ℓ*-TASEP, we found that the limiting occupation density *ρ*(*x, t*) := ℙ(*τ_k_*(*t*) = 1), where 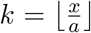, satisfies the nonlinear PDE

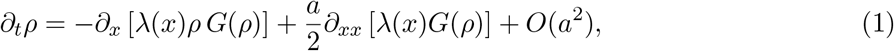

where 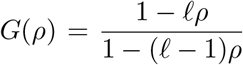 and *λ* is a differentiable extension of (*p*_1_, …, *p*_*N*_), such that *λ*(*x*) = *λ*(*ka*) = *p_k_*. More generally, this PDE takes the form of a conservation law with systematic and diffusive currents *J* and *J_D_*, given by

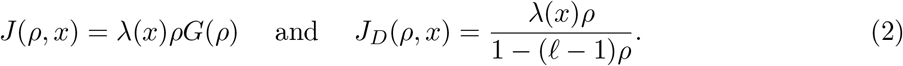

As *a* ≪ 1, the systematic current dominates and solutions of (1) generically converge locally uniformly on (0, 1) to so-called entropy solutions of

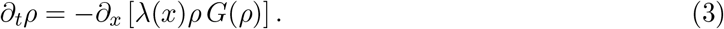

Further details and relevant calculations are provided in **Materials and Methods** and Supplementary Text.

### Particle densities, currents, and phase transitions

The first order nonlinear PDE given by (3) can be solved using the method of characteristics [13], which describes the evolution of differently dense “patches” of particles over time. Solving for the characteristics yields two branches of solutions, which we call “upper” and “lower” branches, while the boundary conditions imposed by *α* and *β* determine which branch is taken by the stationary density of particles (see **Materials and Methods**). As a consequence, the behavior of the system is characterized by a phase diagram in *α* and *β*. Surprisingly, this phase diagram depends on only few parameters of the system (see Figure 2a): the size of particles *ℓ*, the jump rates at the boundaries, *λ*_0_ := *λ*(0) and *λ*_1_ := *λ*(1), and the minimum elongation rate *λ*_min_ := min{*λ*(*x*) : *x* ∈ [0, 1]}. In particular, these parameters determine the critical initiation and termination rates, *α*\* and *β*\*, that are associated with phase transitions. Specifically, *α*\* is given by

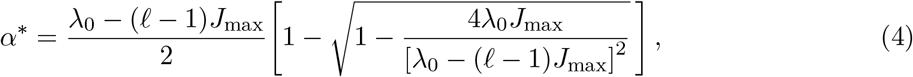

where 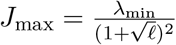, while *β*\* is obtained from (4) by replacing *λ*_0_ with *λ*_1_. The resulting phase diagram, which generalizes previous formulas for the homogeneous 1-TASEP [10], is summarized as follows (see Figure 2b):

1. If *α* < *α*\* and *β* > *β*\* (LD I): In this regime the flux is limited by the initiation rate, leading to a *low density* profile. The corresponding current assumed by the system is

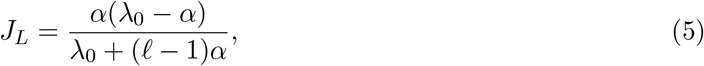

while the site-specific particle density is

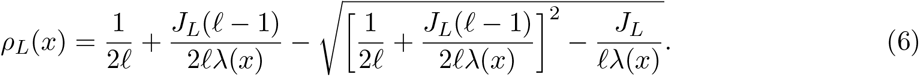
2. If *α* > *α*\* and *β* < *β*\* (HD I): Now the flux is limited by the particle exit rate, resulting in a *high density* regime. The associated current *J_R_* and density *ρ*_*R*_ are identical to *J_L_* (5) and *ρ*_*L*_ (6), respectively, with *λ*_0_ and *α* replaced by *λ*_1_ and *β*.
3. If *α* < *α*\* and *β* < *β*\* (LD II and HD II): The steady state is determined by the sign of *J_L_* - *J_R_* (computed as above). If it is positive (*J_L_* > *J_R_*), the system is in a low density regime with current and density given by *J_L_* and *ρ*_L_, respectively. Conversely, if it is negative, the system is in a high density regime with *J_R_* and *ρ*_R_ as the current and density.
4. If *α* > *α*\* and *β* > *β*\* (MC): The system carries the *maximum possible current* (also referred to as the transport capacity of the system)

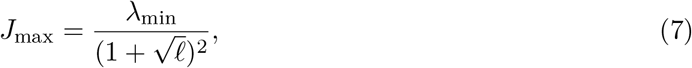

which is limited only by the minimum elongation rate *λ*_min_. Its density is characterized by qualitatively different profiles to the left and right of *x*_min_ = arg min_*x*_ *λ*(*x*): For *x* < *x*_min_, *ρ*(*x*) is described by the upper branch (obtained by replacing *J_R_* with *J*_max_ in the equation for *ρ*_*R*_), while for *x* > *x*_min_, *ρ*(*x*) is described by the lower branch (obtained by replacing *J_L_* with *J*_max_ in *ρ*_L_). That is, a branch switch occurs at *x*_min_ (where 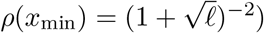). We proved more generally that every global minimum of *λ* regulates the traffic of particles (like a toll reducing the traffic flow) in this fashion: incoming densities to the left of it are always described by the upper branch whereas outgoing particles on the right follow the lower branch. Interestingly, this implies that in the case of multiple global minima, the density between two consecutive minima must undergo a discontinuous jump from lower to upper branch (for more details, see Supplementary Text)

**Figure 2.**
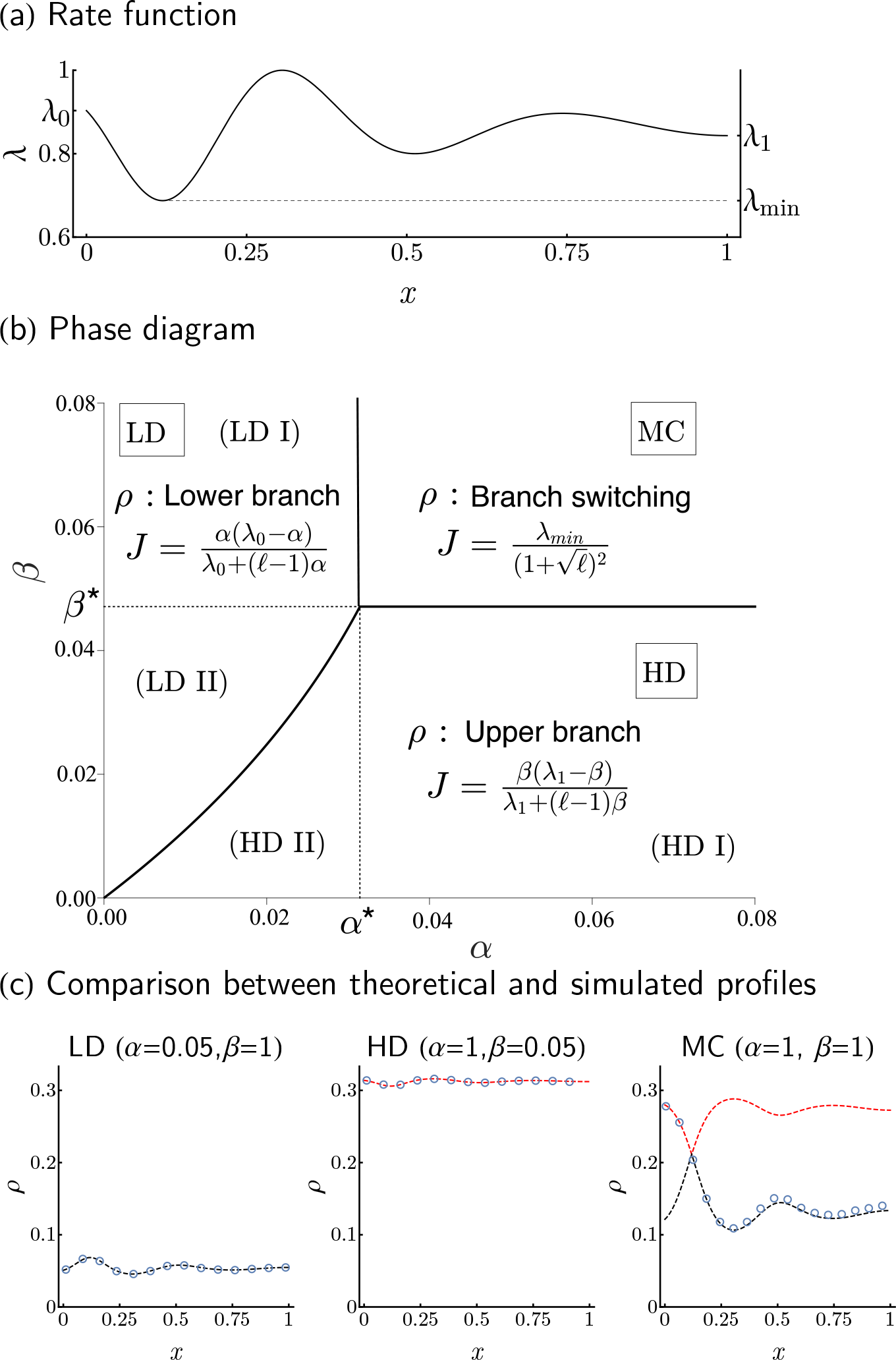
Phase diagram. **(a)** Example rate function with key parameters shown. **(b)** The phase diagram is completely determined by *λ*_0_, *λ*_1_, *λ*_min_ and *ℓ*. In this example, (*λ*_0_, *λ*_1_, *λ*_min_, *ℓ*) = (0.9, 0.3, 0.1, 10). All phase transitions are continuous in *J* and, unless *λ*_min_ coincides with *λ*_0_ or *λ*_1_, discontinuous in **ρ**. **(c)** Simulated results for *ℓ* = 3, *N* = 800, and *λ* as in (a) are compared with theoretical predictions. Dashed black and red lines represent upper and lower branches of solutions to (1). Circles are averaged counts over 5 × 10^7^ Monte-Carlo steps after 10^7^ burn-in cycles.

### Novel phenomena

As shown in Figure 2c, the densities predicted by our analysis agree well with Monte Carlo simulations in all regimes of the phase diagram. We highlight a few novel phenomena in our generalization: First, extending particles to size *ℓ* > 1 and lowering the limiting jump rate *λ*_min_ reduces both the transport capacity *J*_max_ and the critical rates (*α*\* and *β*\*) for entrance and exit, leading to an enlarged MC phase region. This is expected as fewer particles are needed to saturate the lattice, and distances between particles are larger, which in turn limits the number of particles able to cross a site per given time. This phenomenon is quantified precisely using our explicit expressions for *α*\*, *β*\*, and *J*_max_ (see (4) and (7)). Second, the inhomogeneity in *λ* may deform the LD-HD phase separation from being a straight line in the homogeneous *ℓ*-TASEP [14] to a generally nonlinear curve (see Figure 2b) determined by solutions (*α*, *β*) of

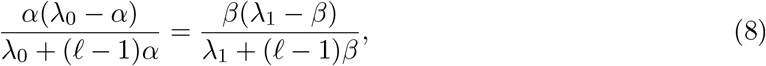

corresponding to the condition *J_L_* = *J_R_*. This is a consequence of *α* and *β* affecting the system at different scales whenever *λ*_0_ ≠ *λ*_1_, resulting in a phase diagram that is no longer symmetric. Lastly, our observation of density profiles performing branch switching in the MC phase was indiscernible in the homogeneous case, as the high density and low density branches merge into a single value (viz. 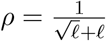).

## 3 Design principles for translational systems

We sought to apply our theoretical analysis to understand how the translational system can be regulated and optimized with regards to protein synthesis rate and ribosome usage. The hydrodynamic theory developed above singles out the key parameters that determine the current and particle densities. We illustrate in Figure 3 how *λ*_0_, *λ*_min_, and *x*_min_ impact the current capacity, its sensitivity to the initiation rate *α*, and the global particle density, suggesting the following principles:

**Figure 3.**
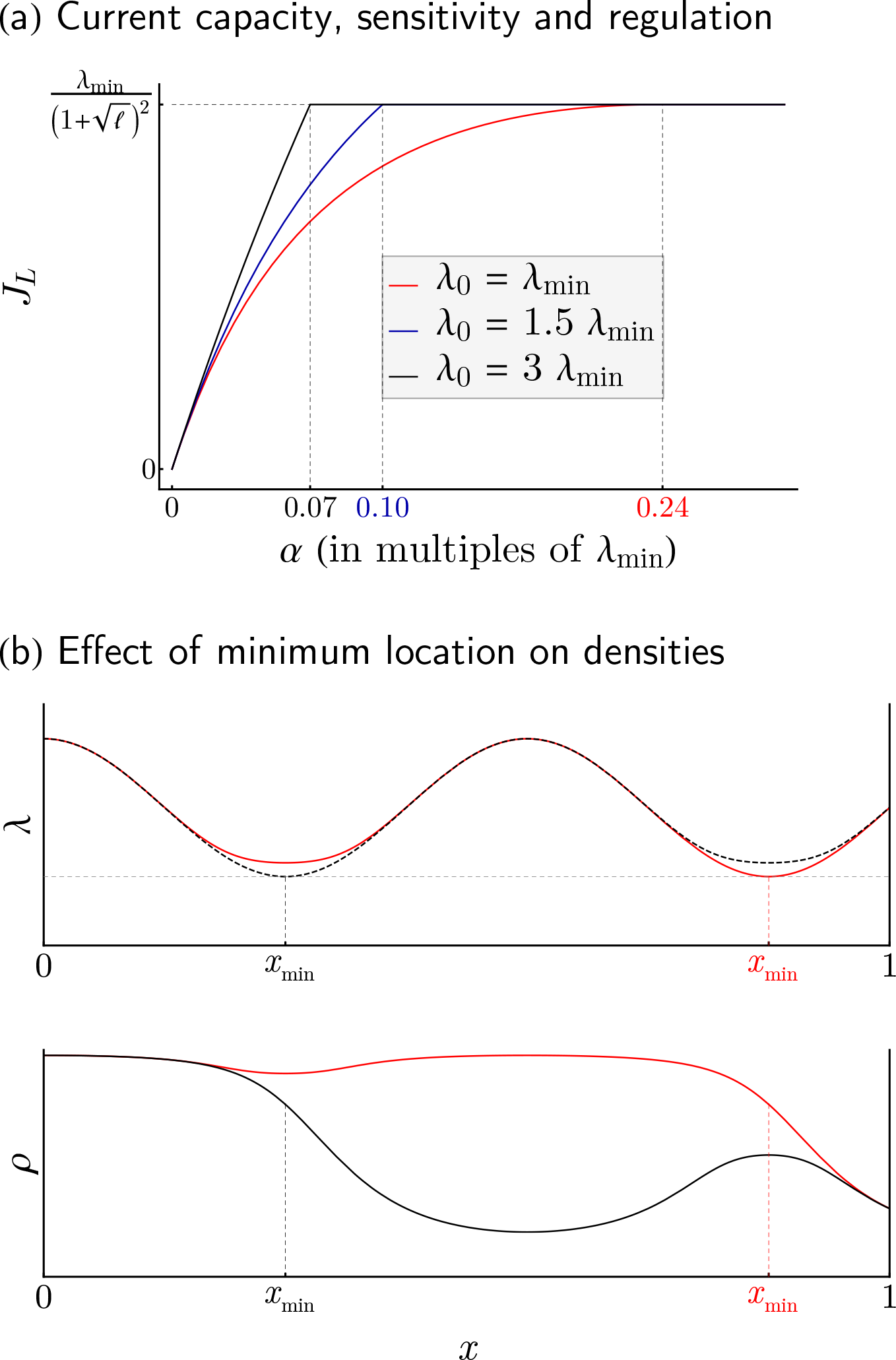
Main determinants of current and particle densities. **(a)** We plot the current *J_L_* in LD/MC against initiation rates *α* for various choices of *λ*_0_. While *λ*_min_ governs the maximum current at which *J_L_* reaches a plateau (coinciding with the transition from LD to MC), changing the size of *λ*_0_ results in changes in ∂_*α*_*J_L_*, the sensitivity of *J_L_* with respect to *α*. Distinct configurations of *λ*_min_ and *λ*_0_ give rise to vastly different dependencies of *J_L_* on *α*, suggesting different responses to global changes in the ribosome pool. The numbers 0.07, 0.10, and 0.24 correspond to *α*\* values in units of *λ*_min_. **(b)** Two elongation rate profiles that differ slightly in overall shape, but drastically in their position *x*_min_ of minimum elongation are plotted (top panel) together with their associated MC ribosome densities (bottom panel). The branch switching phenomenon has extreme consequences for equilibrium particle densities and hence ribosomal costs, with elongation profiles achieving minimum rates close to the initiation site (top, dotted black curve) benefiting from drastic savings (bottom, black curve) compared to otherwise similar profiles (red curves).

### 1. The initiation rate α (and not termination rate β) should regulate the production rate J

As shown by our analysis of the current, any value of the current that lies below the system’s production capacity *J*_max_ can be attained through either HD or LD regime. In order to avoid overuse of resources, however, a transcript should always operate in LD, where the main determinant for currents is the initiation rate *α* (cf. (5)). To guarantee LD profiles, termination rates merely need to exceed the critical value *β*\*, whereas initiation rates are more tightly controlled, varying between 0 and *α*\*. Within this interval, the current *J* increases with *α* according to (5), as illustrated in Figure 3a.

### 2. The minimum elongation rate *λ*_min_ determines the production capacity J_max_

As *α* increases in the LD regime, the current *J* reaches a plateau that is associated with the maximal current (MC) regime (see Figure 3a). By (7), the maximum possible current is directly proportional to *λ*_min_, which therefore sets the range within which production rates may vary. Large values of *λ*_min_ allow for both constitutively high expression of genes as well as highly variable protein levels, while small values of *λ*_min_ guarantee constitutively low expression.

### 3. The sensitivity of production rate to α is moderated by *λ*_0_ and varies across different values of α

As shown in (5), the degree to which *J* varies with *α* is fully determined by the elongation rate *λ*_0_. Indeed, *λ*_0_ controls the time spent by particles at the start of the lattice, and can induce significant buffering if *α* is large enough, thereby modulating the effective rate of entrance associated with *J*. We illustrate this in Figure 3a, where we compare how the current varies as a function of *α* for different values of *λ*_0_ relative to *λ*_min_. The figure also shows that for *λ*_0_ fixed, the flux of a system closer to the MC regime (i.e., with *α* just below *α*\*) is less sensitive to changes in *α*. More generally, the *α*-sensitivity of *J* increases as *λ*_0_ increases. While the variation of the current in *α* is sublinear for *λ*_0_ = *λ*_min_, it becomes linear as *λ*_0_ gets large (see (5)). This suggests in particular that changes in the free ribosome pool (changing the initiation rate globally) can impact the protein production rate differently across different genes.

### 4. Positioning *λ*_min_ close to the start site can reduce the amount of ribosomes used

At maximum production capacity (MC regime), we have shown that the density profile follows the high density branch from the start of the lattice until the location *x*_min_ of *λ*_min_ whereafter it adopts the low density branch. This characteristic branch switching phenomenon makes *x*_min_ critical for the purpose of resource allocation. In Figure 3b, we illustrate how a small local change in the rate function can induce a large increase of average particle density when *x*_min_ changes substantially. Therefore, a way to limit the excessive usage of ribosomes induced by traffic jams at maximum capacity is to position the minimum rate close to the start. However, as previously shown, positioning it too close to the start (such that *λ*_0_ = *λ*_min_) would also decrease the sensitivity of the system to *α*.

## 4 Translational efficiency in yeast

In light of the aforementioned principles, we explored the extent to which the translational system in yeast is efficient. For this study, we used the rates previously inferred from ribosome profiling data for a set of 850 genes in *S. Cerevisiae* [15] (see **Materials and Methods**). We analyzed the location of these genes in the phase diagram, and the distribution of the key parameters and variables that determine the ribosomal currents and densities. Interestingly, we found the theoretical design principles being reflected as follows:

### 1. Translation mainly operates in LD regime

Upon computing *α*\* and *β*\*, we located the position of each gene in the phase diagram (see Figure 4a). Over the 850 genes in our dataset, we found 841 in LD and the remaining 9 in the MC region. No genes were found in HD, suggesting no excessive usage of ribosome to achieve any protein level. As a result, the initiation rate is the main determinant and limiting factor of the current (Spearman’s rank correlation coefficient **ρ** = 0.979). The strength of this correlation nevertheless decreases as genes get closer to the MC regime, since *J* becomes less sensitive to *α* and *λ*_min_ becomes its rate limiting factor (see Figure S6a). To quantify this reduction in correlation, we binned the data by quartiles of *J* and computed Spearman correlations within each bin, which yielded (in order of quartiles): 0.93, 0.72, 0.64, and 0.58.

**Figure 4.**
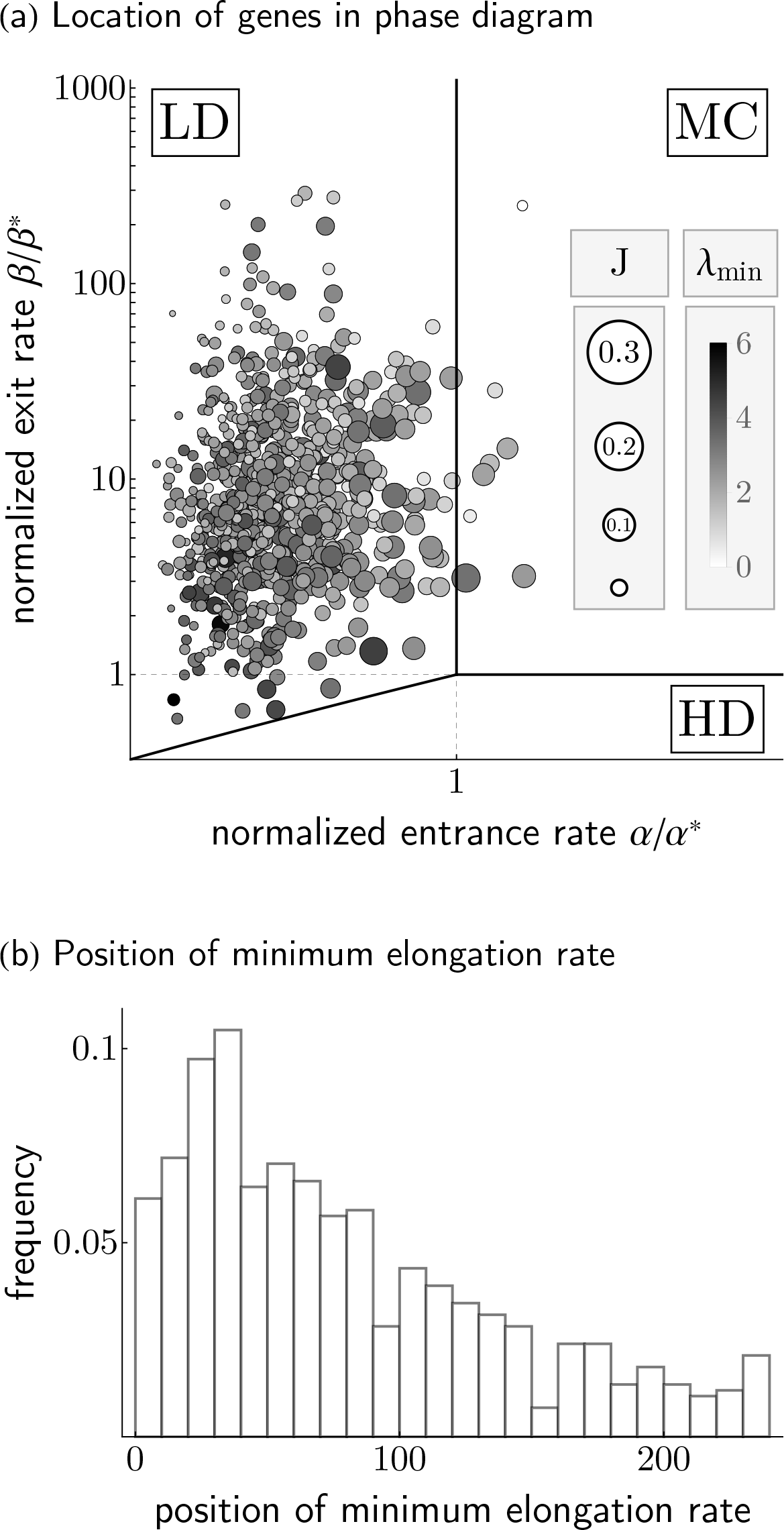
Translation machinery in *S. cerevisiae* optimizes for ribosomal cost, flexible regulation and production capacity. **(a)** 850 genes of *S. cerevisiae* are located in the phase diagram, with size and hue of each data point reflecting current and minimum elongation rate, respectively. On a population level, systems of comparable production capacities (∝ *λ*_min_) fully exploit their dynamic range by adjustment of *α*, with highly expressed proteins likely situated inside or close to MC. **(b)** The resulting resource cost considerations drive a significant number of transcripts to position their minimum elongation rate early on in the codon sequence, forcing ribosomal traffic jams to remain short.

### 2. Wide ranges of currents are covered within production capacity

For each gene in our dataset, we examined the maximal protein production rate, which according to our theory is proportional to *λ*_min_. The data exhibit an overall range of *λ*_min_ between 1.01 and 6.01 codons/second, and for any fixed *λ*_min_, currents are well spread out across [0, *J*_max_] (see Figure S6b). Given that genes cover almost all of the theoretically possible range of currents, we investigated whether certain configurations of *λ*_min_ and *J* are associated with the biological function of specific genes. To do so, we compared ribosomal protein genes (known to be highly expressed) and genes related to stress response (requiring variable expression over time, see **Materials and Methods**). We found that, while both sets of genes display comparable *λ*_min_, ribosomal genes are more likely to be close to their maximal production capacity (*p* < 7 × 10^−3^, see **Materials and Methods**) and more consistently so (the coefficient of variation is 0.22 for ribosomal genes and 0.36 for stress response).

### 3. *λ*_0_ (associated with sensitivity to α) is higher for genes that are either highly expressed or subject to varying expression demand

The impact of increasing *α*-sensitivity is primarily twofold: First, for fixed production capacity, large currents may be attained with smaller initiation rates; and second, more substantial changes in currents may be achieved with small changes in *α*. To investigate the former we computed *α*\*, the critical rate necessary for a gene to attain maximum capacity, across all genes whose *λ*_min_ exceeded the median *λ*_min_ of the data set (as large currents presuppose large capacities). Further binning this range into quartiles (to isolate the dependence of *α*\* on *λ*_0_), we found that genes whose currents are at least 90% of the production capacity are significantly more sensitive (*p* < 0.008, 0.01, 0.05, and 0.004, respectively; see Figure S6c), requiring smaller initiation rates to reach peak production rate (cf. Figure S6a). To inspect the second aspect of *λ*_0_ as facilitator or inhibitor of rapid changes in current, we explored the ratio of *λ*_0_ to *λ*_min_ again in ribosomal and stress response genes. For constitutively highly expressed genes like ribosomal genes, we expect this ratio to be small to maintain stable current close to MC (cf. Figure 3), whereas genes with variable expression demands like the ones associated with stress response should exhibit larger ratios. Confirming this intuition, we found significantly reduced levels of *λ*_0_/*λ*_min_ in ribosomal genes (*p* < 2 × 10^−6^), and significantly increased levels in stress response genes (*p* < 0.04).

### 4. The position of *λ*_min_ is preferentially located early in the open reading frame

Upon analyzing the distribution of *x*_min_ from our dataset (see Figure 4b), we found it preferentially located in the codon positions between 30 to 40, consistent with genes forestalling excessive ribosome usage through enforcing branch switching early on. More specifically, we reasoned that both genes closer to MC and those highly sensitive to *α* run higher risk of incurring substantial ribosome cost and should thus locate *x*_min_ early in the coding sequence. Indeed, both the top quartile of genes close to MC (as measured by *α/α*\*) and stress response associated genes showed significantly smaller *x*_min_ (*p* < 0.03 and 0.01, respectively). Moreover, genes with unusually large values of *x*_min_ are significantly less likely to be close to MC (top quartile of *x*_min_: *p* < 1 × 10^−3^).

## 5 Discussion

While past quantitative studies of the TASEP under general conditions of extended particle size and/or rate heterogeneity have mostly been limited to numerical simulations or mean-field approximations [14, 16, 17, 18, 19], we used here a different approach that relies on studying the hydrodynamic limit of the process. In the case of homogeneous rates, previous studies [11, 20] established this hydrodynamic limit, but without further analyzing the subsequent PDE. After deriving this limit for inhomogeneous rates, we obtained closed-form formulas for the associated current, densities, and phase diagram, generalizing previous theoretical results for the TASEP [9, 10] and its variants [14, 17, 21]. Our approach has the advantage of revealing the key parameters that the current and densities depend on, enabling an immediate quantification of the process and its phase diagram. Such a quantification is difficult to achieve via conventional stochastic simulations or approximations used in the past several years [7, 8, 22].

Our characterization of the current and densities in the phase diagram suggests that, in agreement with earlier experimental studies [23, 24], translation dynamics should be mainly governed by the initiation rate, while the termination rate and most of elongation rates have negligible impact. In particular, our results explain why having the initiation rate as the main limiting factor of the current [25] minimizes ribosome usage. In addition, we identify two critical parameters of the system, namely, the elongation rate immediately following initiation and the minimal elongation rate. Previous studies have established some association between the sequence context in the early 5′ coding region and protein production levels [26, 27]. For example, it has been shown that mRNA secondary structure in the first ~ 16 codons (which locally decreases the elongation rate) negatively affects the translation rate in *E. Coli*, while no significant contribution of mRNA folding in other regions was found [27]. By exposing *α* and *λ*_0_ as the only parameters that currents in LD depend on, our analysis suggests a direct explanation for such contrast.

We also highlighted the impact of *λ*_0_ on the sensitivity of the current to changes in *α*. In practice, initiation rates can vary at the individual gene level (e.g., through interactions with specific miRNAs [28]). According to our theory, the way that these variations impact the protein production rate depends on *λ*_0_; we hence suggest that this may explain why genes associated with stress response present higher values of *λ*_0_, as it facilitates the response to changes in *α*. At a more global level, our study shows how protein levels can be more or less robust against changes in the ribosomal pool, which can simultaneously affect all initiation rates in a cell. Since the level of ribosomes present in a cell fluctuates over time [29], it would be interesting to see if protein levels scale uniformly with these variations across genes, and if not, whether the differences in *λ*_0_ can explain it.

To the best of our knowledge, the role of the minimum elongation rate *λ*_min_ has so far received attention only indirectly, through the study of what is known as the “5′ translational ramp” [30]. This ramp is a pattern of translational slowdown around codon position 30-50 followed by steadily accelerating elongation rates, which is mirrored by the spatial distribution of minimum elongation rates we found here. This ramp has been hypothesized to prevent crowding of ribosomes on the transcript [30], for which we provide a theoretical basis, exposing *λ*_min_ as a separator between crowded and freely elongating ribosomes. More generally, the complex interplay between the maximum current capacity, ribosome usage, and sensitivity to the initiation rate suggests various ways to set the parameters *λ*_0_, *λ*_min_ and *x*_min_, depending on the desired object to optimize. For example, allocating the minimum elongation rate near the beginning of the ramp region provides an optimal trade-off between high sensitivity and minimal traffic jams. On the other hand, it would be optimal for genes with housekeeping function to have a decreased sensitivity, which would push the minimum to earlier positions.

Finally, our analysis can help to answer the long-debated question regarding the implication of translation on codon usage bias [31, 32]. Since highly expressed genes are enriched for synonymous codons translated by more abundant tRNAs [4, 33], it has been hypothesized that codon usage bias increases translation efficiency by accelerating elongation [31]. However, recent studies have challenged such a hypothesis, suggesting that translational selection for speed is not sufficient to explain the observed variation in codon usage bias [34]. Synonymous changes of the coding sequence modify local elongation rates, but, according to our theory, such a modification impact the overall protein production rate only if *λ*_0_ or *λ*_min_ is affected. In addition, our work implies that synonymous codon replacements that substantially change the location *x*_min_ of *λ*_min_ affect the efficiency of ribosome usage, and hence are more likely to be under selective pressure. Aside from these cases, there should be little direct impact of synonymous codon usage on translation efficiency; this prediction is consistent with previous studies that attempted to explain differences in expression using codon identity [35]. Other factors such as mRNA decay [4], or reduction of nonsense errors or co-translational misfolding [32, 36] might be more important drivers of codon usage bias.

## 6 Material and Methods

### Analysis of the conservation law using the methods of characteristics

The conservation law in (3) is solved by using the methods of characteristics. The characteristics are solutions of the system of ordinary differential equations

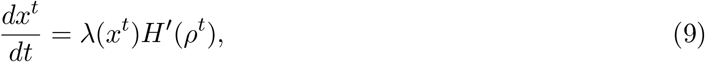

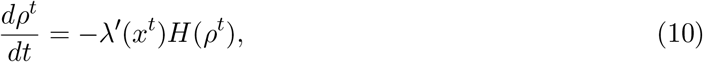

where *H*(**ρ**) = **ρ*G*(**ρ**), and *H*′ and *λ*′ denote the derivatives of *H* and *λ*, respectively. Initial information is propagated across space-time through these characteristics. Studying the stationary solutions of (3) thus requires evaluating the boundary densities imposed by *α* and *β*, and determining which of these dominates the lattice at equilibrium (see Supplementary Text).

### Statistical tests and computation of *p*-values

To establish significance of a subset *X* of genes with respect to a statistic *f* (e.g., *α, J* or *x*_min_) relative to a background set *Y*, we performed hypothesis testing on the median *m_f_* of *f* over samples in *X*. Under the null distribution of *X* being drawn uniformly at random, the probability of this test statistic exceeding *m* equals the probability of a hypergeometric variable with parameters *N* = |*Y*|, *K* = 2 |*Y_m_*|, *n* = |*X*|, where *Y_m_* is the set of genes in *Y* whose *f* exceeds *m*, exceeding ⌊|*X/*2|⌋. This p-value can be computed explicitly. Sets of ribosomal and stress response genes were taken from the Saccharomyces Genome Database [37].

### Data processing

Initiation, elongation, and termination rates were taken from an earlier work [15], where the rates were estimated from ribosome profiling data of *S. Cerecisiae* for a set of 850 genes selected based on length and footprint coverage. The *α* and *β* values were taken directly from that publication. To compute the elongation rates corresponding to the hydrodynamic limit, which treats densities over *ℓ* sites (see Supplementary Text), we applied a ten-codon moving average to the elongation rates from [15].

## Acknowledgments

This research is supported in part by an NIH grant R01-GM094402, and a Packard Fellowship for Science and Engineering. YSS is a Chan Zuckerberg Biohub Investigator.

